# A Systems-Level Model Reveals That 1,2-Propanediol Utilization Microcompartments Enhance Pathway Flux Through Intermediate Sequestration

**DOI:** 10.1101/069542

**Authors:** Christopher M. Jakobson, Marilyn F. Slininger, Danielle Tullman-Ercek, Niall M. Mangan

## Abstract

The spatial organization of metabolism is common to all domains of life. Enteric and other bacteria use subcellular organelles known as bacterial microcompartments to spatially organize the metabolism of pathogenicity-relevant carbon sources, such as 1,2-propanediol. The organelles are thought to sequester a private cofactor pool, minimize the effects of toxic intermediates, and enhance flux through the encapsulated metabolic pathways. We develop a mathematical model of the function of the 1,2-propanediol utilization microcompartment of *Salmonella enterica* and use it to analyze the function of the microcompartment organelles in detail. Our model makes accurate predictions of doubling times based on an optimized compartment shell permeability determined by maximizing metabolic flux in the model. The compartments function primarily to decouple cytosolic intermediate concentrations from the concentrations in the microcompartment, allowing significant enhancement in pathway flux by the generation of large concentration gradients across the microcompartment shell. We find that selective permeability of the microcompartment shell is not absolutely necessary, but is often beneficial in establishing this intermediate-trapping function. Our findings also implicate active transport of the 1,2-propanediol substrate under conditions of low external substrate concentration, and we present a mathematical bound, in terms of external 1,2-proanediol substrate concentration and diffusive rates, on when active transport of the substrate is advantageous. By allowing us to predict experimentally inaccessible aspects of microcompartment function, such as intra-microcompartment metabolite concentrations, our model presents avenues for future research and underscores the importance of carefully considering changes in external metabolite concentrations and other conditions during batch cultures. Our results also suggest that the encapsulation of heterologous pathways in bacterial microcompartments might yield significant benefits for pathway flux, as well as for toxicity mitigation.

**Author Summary:** Many bacterial species, such as *Salmonella enterica* (responsible for over 1 million illnesses per year in the United States) and *Yersinia pestis* (the causative agent of bubonic plague), have a suite of unique metabolic capabilities allowing them to proliferate in the hostile environment of the host gut. Bacterial microcompartments are the subcellular organelles that contain the enzymes responsible for these special metabolic pathways. In this study, we use a mathematical model to explore the possible reasons why *Salmonella* enclose the 1,2-propanediol utilization metabolic pathway within these sophisticated organelle structures. Using our model, we can examine experimentally inaccessible aspects of the system and make predictions to be tested in future experiments. The metabolic benefits that bacteria gain from the microcompartment system may also prove helpful in enhancing bacterial production of fuels, pharmaceuticals, and specialty chemicals.

## Introduction

Bacterial microcompartments (MCPs) are protein-bound intracellular organelles used by *Salmonella enterica*, *Yersinia pestis*, *Klebsiella* spp., and other bacteria to spatially organize their metabolism [1–3]. MCP metabolons allow the growth of these pathogens on carbon and energy sources, such as 1,2-propanediol [4] and ethanolamine [5], that confer a competitive advantage upon invasion of the host gut [6–10]. MCPs are typically approximately 150 nm in diameter, with multiple enzymes localized inside a porous, monolayer shell composed of several distinct proteins [4,11,12]; a typical bacterial cell contains several MCP structures when in the presence of the appropriate substrate. Enzymes are localized to the MCP interior through the interactions of N-terminal signal sequences with the inward-facing helices of MCP shell proteins, and potentially through other uncharacterized interactions [13–15].

Inside the 1,2-propanediol utilization (Pdu) MCP metabolon, 1,2-propanediol metabolism proceeds as follows: the vitamin B12-dependent PduCDE holoenzyme converts 1,2-propanediol to propionaldehyde [14], then propionaldehyde is converted to either 1-propanol by the NADH-dependent PduQ enzyme [16] or to propionyl-coA by the NAD+-dependent PduP enzyme [17] (Fig. 1A). The PduP and PduQ enzymes are thought to cycle a private pool of NAD+ /NADH inside the MCP lumen, enforcing a 1-to-1 stoichiometry for the two reactions [18,16,19]. 1-propanol is not used for cell growth, but propionyl-CoA can be utilized either as a carbon source or for ATP generation through substrate-level phosphorylation [20].

**Figure 1.**
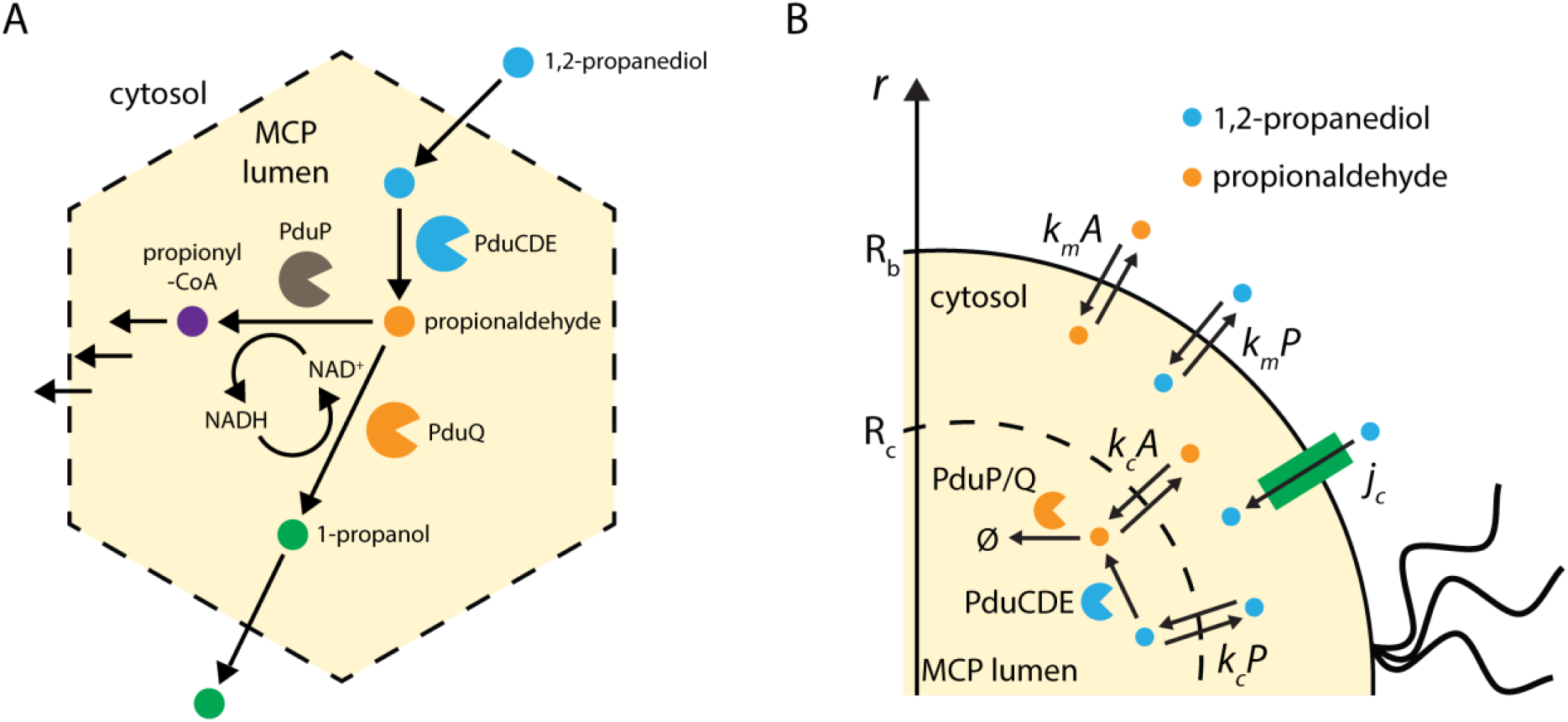
Reaction scheme for (A) the native Pdu MCP and (B) the simplified model considered here.

Pdu MCPs are elaborate multi-protein structures subject to exquisite regulation, and much investigation has focused on determining the detailed function of the organelles. Experiments suggest that metabolic pathways are sequestered in the Pdu and ethanolamine utilization (Eut) MCPs in order to protect the cell from toxicity associated with aldehyde intermediates [21], to prevent carbon loss from the metabolic pathway [19], and to provide a private pool of cofactors for the encapsulated pathways [18,19]. These mechanistic hypotheses are difficult to confirm experimentally, as directly measuring the concentrations of small molecules inside the MCPs *in vivo* remains a challenge. Here we build a coupled reaction-diffusion model of the Pdu MCP and use computational and analytic approaches to assess whether the described biological system produces the hypothesized mechanistic behavior. The use of a mechanistic model of the MCP allows us to examine potential functions and behavior across a wide range of parameters, providing a framework to incorporate emerging experimental observations and guide the design of future experiments.

The model presented here follows an approach used to investigate the function of a related organelle, carboxysomes in the carbon concentrating mechanism of cyanobacteria [22]. For simplicity, we model the Pdu MCP as a spherical compartment in the center of a radially symmetric spherical cell. The model includes passive transport of 1,2-PD and propionaldehyde across the cell membranes and MCP shell, possible active transport of 1,2-PD into the cell, and the action of the PduCDE and PduP/Q enzymes localized within the MCP (Fig. 1B). Parameters were estimated *a priori* or based on experimental results. We have developed a numerical simulation for this spherical geometry with localization of enzymes to the MCP. By making the assumption of constant metabolite concentrations in the MCP lumen, we also developed a closed-form analytic solution that well approximates the full numerical solution for a broad range of physically relevant parameter values (Fig. S1; see also Models). The analytic approximation allows for explicit examination of the relationships between different parameters and the mechanisms in the system. This analytical solution is used throughout the following analysis.

We find that aldehyde sequestration is the key function of the Pdu MCP, and contributes not only to decreasing aldehyde leakage into the cytosol and the growth medium, as is often discussed in the existing literature, but also to greatly increasing flux through the metabolon by increasing the substrate concentration in the vicinity of the relevant enzymes. Furthermore, we find that active 1,2-PD transport across the cell membrane is dispensable at some external 1,2-PD concentrations, including the concentrations at which most laboratory experiments are performed, but not at low external 1,2-PD concentrations. This transport activity has been proposed previously, but never experimentally observed [25]. Finally, while selective MCP membrane permeability is not always required to achieve optimal substrate concentrations, it is often advantageous in this regard. The qualitative behavior of our model and quantitative fluxes and metabolite concentrations agree well with existing experimental results, without fitting any model parameters to experimental data. Additionally, our results suggest several avenues for continuing computational and experimental investigation, including investigations into 1,2-PD active transport, direct characterization and detailed simulation of MCP membrane permeability, and analysis of MCP function in chemostatic cultures.

## Models

We model Pdu MCP function using a simple spatially resolved reaction-diffusion model incorporating passive and active transport across the cell membrane, passive transport across the MCP shell, and enzymatic catalysis of two critical steps in 1,2-propanediol metabolism: conversion of 1,2-propanediol to propionaldehyde by the PduCDE holoenzyme [9], and subsequent conversion of propionaldehyde to downstream products by the PduP and PduQ enzymes [11,12]. We model the bacterial cell as a spherical compartment of radius 500 nm, containing at its center a single spherical MCP of radius 100 nm (representing the same fraction of the cell volume as 5 Pdu MCPs in a typical bacterial cell; Fig. 1B).

The key assumptions in the model are as follows:

1. we assume that the cell and the MCP are spherically symmetrical, such that 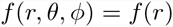 only, and 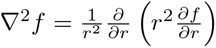
2. we consider the system at steady state;
3. we consider constant external concentrations of 1,2-propanediol (*P*_*out*_) and propionaldehyde (*A*_*out*_);
4. and we assume that the enzyme-catalyzed reactions are irreversible, and neglect reactions downstream of PduP/Q.

We assume that the conversion of *P* to *A* in the absence of enzymatic catalysis is negligible, so the equations for diffusion of 1,2-PD, *P*, and propionaldehyde, *A*, in the cytosol are as follows:

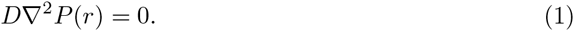

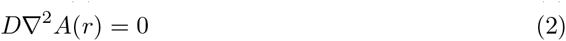

Where *D* is the diffusion coefficient of the metabolites in the cytosol. The analogous equations in the MCP are likewise:

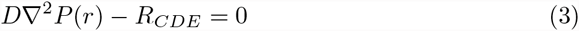

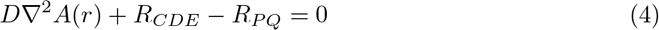

Inside the Pdu MCP, we assume Michaelis-Menten kinetic behavior of the PduCDE and PduP/PduQ enzymes, so the equation for the rate of the PduCDE (diol dehydratase) reaction is

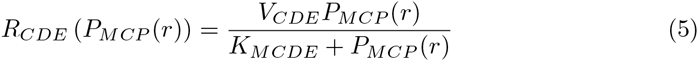

Here *V_CDE_* is the maximum rate of dehydration by PduCDE. *K_MCDE_* is the half maximum concentration for dehydration.

PduP and PduQ are redox-coupled by the cycling of NAD+/NADH, so we assume that their catalytic rates are equal at steady state; the equation for the PduP and PduQ reactions is therefore:

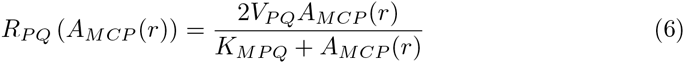

Here *V*_*PQ*_ is the maximum rate of aldehyde consumption by PduP, and *K*_*MPQ*_ is the half maximum concentration. The rate is doubled due to cofactor cycling to yield the rate of PduP/Q combined.

We assume that *P* and *A* are transported across the cell membranes by passive diffusion, so we can specify the following boundary conditions enforcing continuity of flux of each metabolite at the cell membrane. In the case of *P*, in addition to passive transport across the cell membrane, we also include the possibility of active transport across the cell membrane by the putative membrane protein encoded by *pduF*.

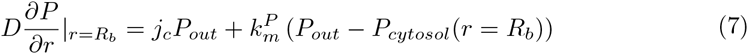

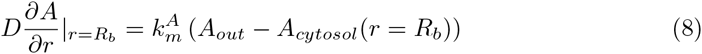

Here active transport of *P* is set by the transport velocity *j*_*c*_. The permeabilities of the cell membrane to *A* and *P* are set by the passive transport velocities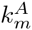 and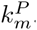

The passive transport velocities of *P* and *A* across the MCP shell can be treated independently or as being equal; we explore the necessity of selective permeability by allowing the velocities to differ. The continuity of flux at the MCP shell sets the linking boundary condition between the concentrations inside the MCP and in the cytosol as follows:

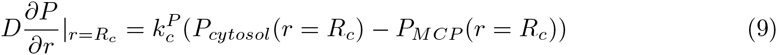

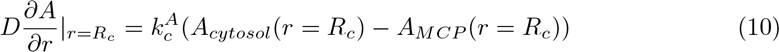

The passive transport, active transport, and enzyme parameters were estimated *a priori* or from literature as shown in Table 1. The nonspecific permeability of the MCP *k*_*c*_ was chosen to minimize the predicted *S. enterica* doubling time (Fig. 2A).

**Figure 2.**
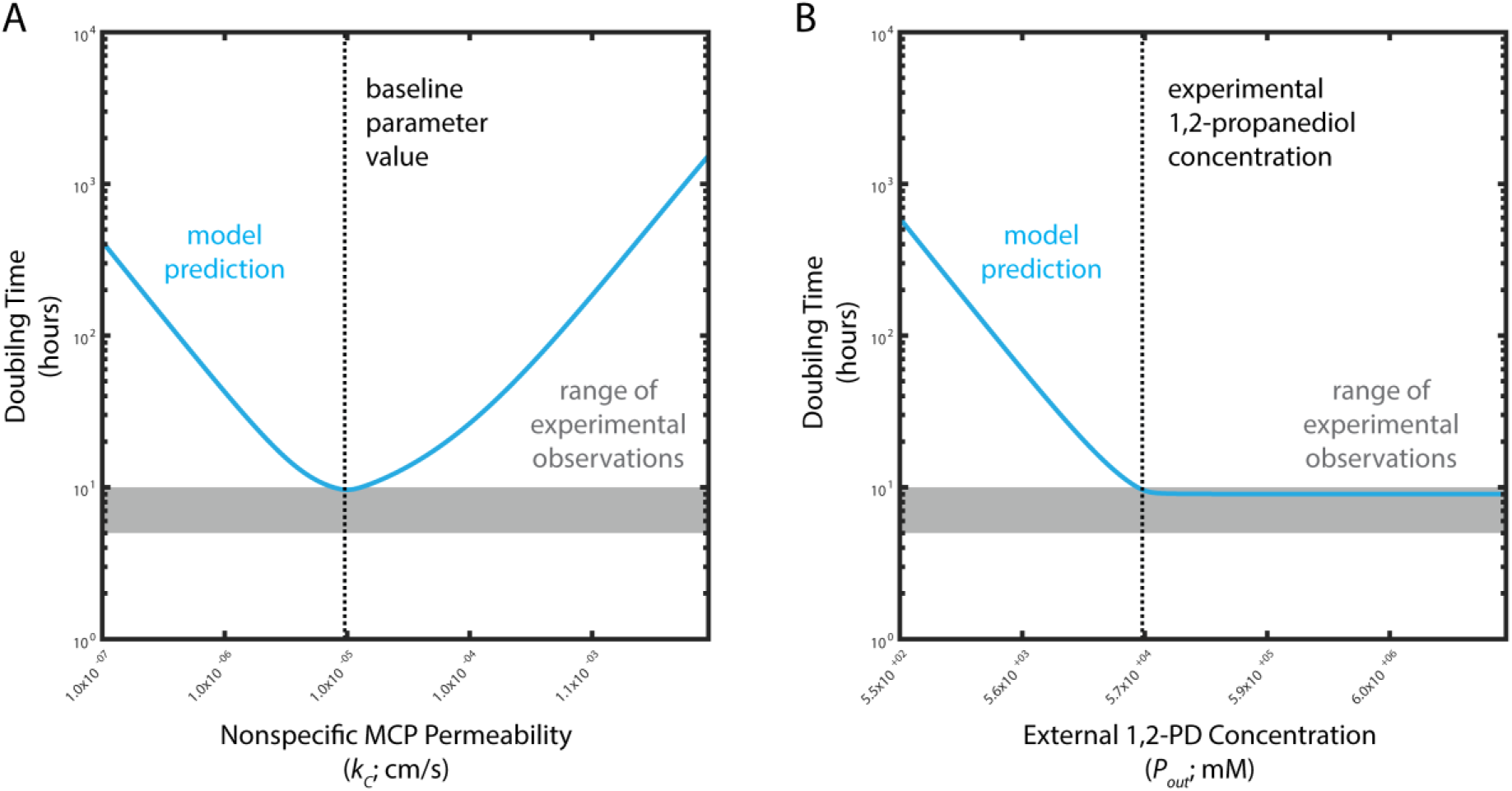
Predicted *S. enterica* doubling times for cells with MCPs as a function of (A) nonspecific MCP permeability k_c_ and (B) external 1,2-PD concentration *P*_*out*_. Model predictions are shown in blue; observed doubling times for experiment are shown by the grey shaded area. The baseline *k*_*c*_ value and the typical experimental external 1,2-PD concentration are shown by black dashed lines in (A) and (B), respectively.

**Table 1.** Parameter estimates used in our model.

| Parameter | Meaning | Estimated Value | Units |
| --- | --- | --- | --- |
| $j_c$ | Rate of active transport of 1,2-PD across the cell membrane | 1 [22] | $\frac{cm}{s}$ |
| $k_c^A$ | Permeability of the Pdu MCP to propionaldehyde | $10^{-5}$ [22] | $\frac{cm}{s}$ |
| $k_c^P$ | Permeability of the Pdu MCP to 1,2-PD | $10^{-5}$ [22] | $\frac{cm}{s}$ |
| $R_b$ | Radius of the bacterial cell | $5 \times 10^{-5}$ | $cm$ |
| $R_c$ | Radius of the Pdu MCP | $10^{-5}$ [22] | $cm$ |
| $D$ | Diffusivity of metabolites in the cellular milieu | $10^{-5}$ [22] | $\frac{cm^2}{s}$ |
| $k_m^A$ | Permeability of the cell membrane to propionaldehyde | $10^{-3}$ [31] | $\frac{cm}{s}$ |
| $k_m^P$ | Permeability of the cell membrane to 1,2-PD | $10^{-3}$ [31] | $\frac{cm}{s}$ |
| $k_{catCDE}$ | Maximum reaction rate of a PduCDE active site | $3 \times 10^2$ [32] | $\frac{1}{s}$ |
| $N_{CDE}$ | Number of PduCDE enzymes per cell | $1.5 \times 10^3$ [12] | $\frac{1}{cell}$ |
| $K_{MCDE}$ | Michaelis-Menten constant of PduCDE | $5 \times 10^2$ [32] | $\mu M$ |
| $k_{catPQ}$ | Maximum reaction rate of a PduPQE active site | 55 [16] | $\frac{1}{s}$ |
| $K_{MPQ}$ | Michaelis-Menten constant of PduPQ | $1.5 \times 10^4$ [16] | $\mu M$ |
| $N_{PQ}$ | Number of PduPQ enzymes per cell | $2.5 \times 10^3$ [12] | $\frac{1}{cell}$ |
| $P_{out}$ | External 1,2-PD concentration | $5.5 \times 10^4$ [21] | $\mu M$ |
| $A_{out}$ | External propionaldehyde concentration | 0 [21] | $\mu M$ |

### Non-dimensional equations

We derive nondimensional equations which can then be solved numerically by a finite-difference approach to find the steady-state concentrations in the MCP, and the solutions in the cytosol follow directly. We solve the spherical finite-difference equations using the `ODE15s` solver in MATLAB. Details of the non-dimensionalization can be found in the Supplementary Information.

### Analytical solution

If we assume that the concentration gradients in the MCP are small, then the concentrations *P*_*M C P*_ and *A*_*M C P*_ are approximately constant and the full solution to the reaction-diffusion equations in the MCP and cytosol can be found analytically. This assumption is tantamount to assuming that the quantity 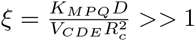 (see Supplementary Information); given our assumptions, we estimate that the value of *ξ* is approximately 10^4^. The detailed solution is shown in Appendix A.

### Equations for no MCP case

In the case when there is no Pdu MCP, we assume that the same number of enzymes are now distributed throughout the cell. We can thence derive nondimensional equations which can be solved numerically by a finite-difference approach as above (see Supplementary Information).

## Results

### MCPs reduce toxicity by decoupling cytosolic aldehyde concentration from PduP/Q saturation

In order to assess the function of the Pdu MCP, we compare the performance of the Pdu MCP system to two alternative organizational strategies for the Pdu metabolic enzymes: uniform distribution of the enzymes throughout the cytosol, and co-localization on a scaffold without a diffusion barrier. We assess the function of each organization strategy by two criteria: (i) maintenance of the cytosolic propionaldehyde concentration below the toxicity limit of 8 mM [21] in Figure 3A, and (ii) saturation of the PduP/Q enzymes with their propionaldehyde substrate in Figure 3B. Flux through the Pdu metabolon is maximized when the enzymes are saturated. In each organizational case, we examine the kinetically relevant propionaldehyde concentration (Fig. 3B): without MCPs, this is the cytosolic propionaldehyde concentration; with MCPs, this is the propionaldehyde concentration in the MCP.

**Figure 3.**
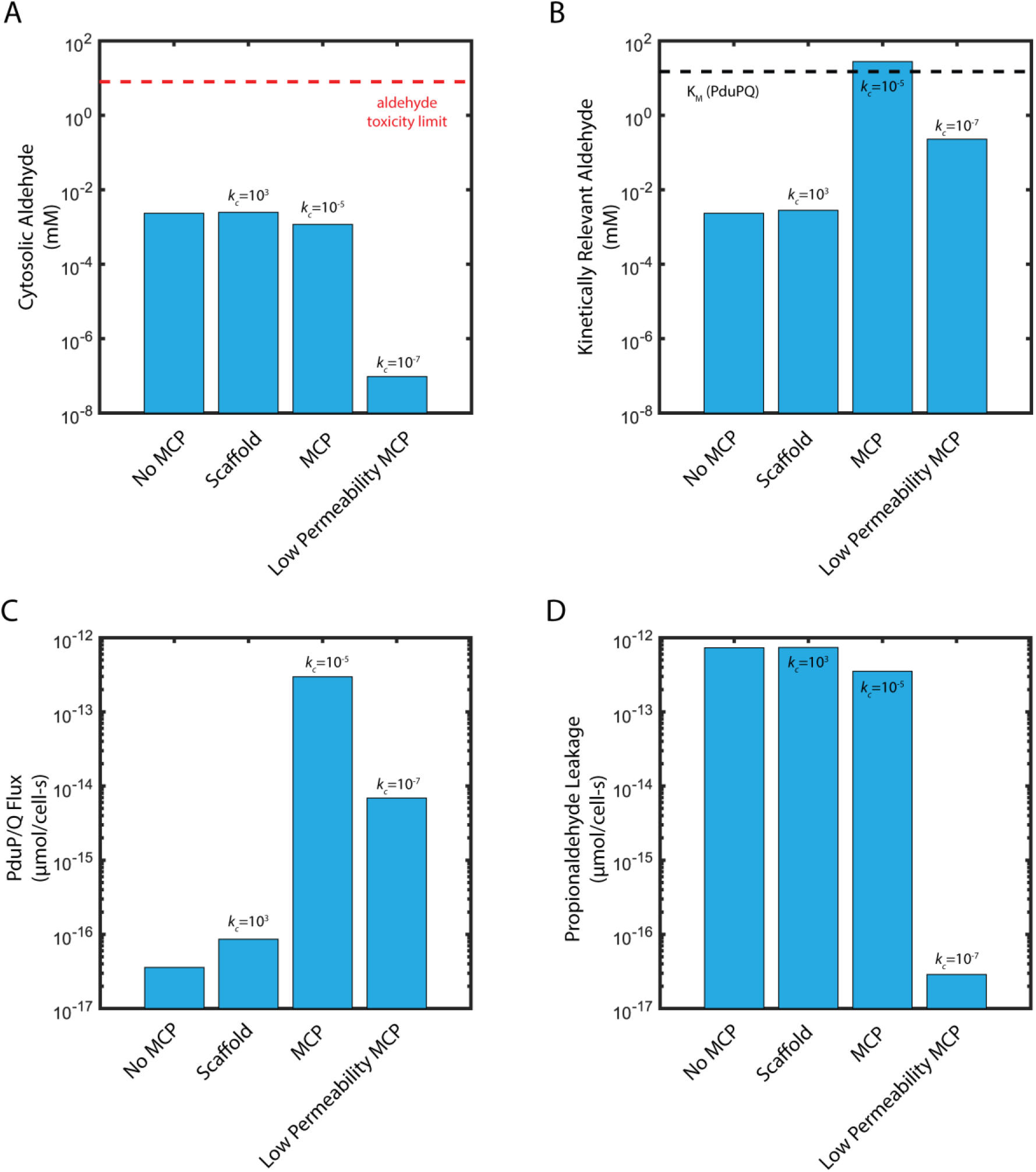
(A) Cytosolic propionaldehyde concentration (*A*_*cyto*_), (B) kinetically relevant propionaldehyde concentration (*A_MCP_* or *A_cyto_*, as appropriate), (C) carbon flux through PduP/Q, and (D) propionaldehyde leakage out of the cell into the extracellular space. Shown are the steady state concentrations and fluxes for cases without MCPs, with a scaffold, with MCPs, and with MCPs of extremely low permeability. The 15 mM KM of PduP/Q is shown as a black dashed line in (B); the 8 mM propionaldehyde cellular toxicity limit is shown as a red dashed line in (A). Baseline parameters are in Table 1.

The case of enzymes distributed throughout the cytosol provides an assessment of the baseline efficacy of the Pdu pathway without compartmentalization. As the Michaelis-Menten constant of the PduP/Q enzymes is approximately 15 mM, above the propionaldehyde toxicity limit, it is impossible to both saturate the PduP/Q enzymes and remain below the toxicity limit in the same location. In fact, if the PduCDE and PduP/Q enzymes are distributed throughout the cytosol, our model suggests that the steady-state propionaldehyde concentration is maintained at 2.4 *μ*M (several orders of magnitude below the 8 mM toxicity limit) when the external propanediol concentration is 55 mM (Fig. 3A, Fig. S2). In turn, the PduP/Q enzymes, with a *K_M_* of 15 mM, are not saturated (Fig. 3B).

Another organizational strategy is localization of the relevant enzymes to a scaffold, without a diffusion barrier. In this case, the propionaldehyde concentration in the vicinity of the PduP/Q enzymes is 2.8 *μ*M, higher than if the enzymes are distributed throughout the cytosol (in which case the kinetically relevant concentration is 2.4 *μ*M), but still much lower than the saturating concentration of 15 mM (Fig. 3B).

When the enzymes are localized in the MCP (permeability of 10^−5^ cm/s for 1,2-PD and propionaldehyde), the PduP/Q enzymes are exposed to a much higher propionaldehyde concentration of 28 mM (higher than the saturating concentration) (Fig. 3B), while the propionaldehyde concentration in the cytosol is 1.2 *μ*M (Fig. 3A). The presence of a diffusion barrier allows the MCP to decouple the cytosolic propionaldehyde concentration (responsible for toxicity) from the kinetically relevant propionaldehyde concentration in the vicinity of the PduP/Q enzymes. Very low nonspecific permeabilities of the diffusion barrier are unfavorable, however: a MCP with very low permeability (10^−7^ cm/s for 1,2-PD and propionaldehyde) maintains a very low cytosolic propionaldehyde concentration of 1 pM, but also a low concentration of propionaldehyde in the MCP of 230 *μ*M (Fig. 3A, B).

Optimally permeable MCPs are an effective means of decoupling a potentially toxic cytosolic aldehyde concentration from the kinetically relevant aldehyde concentration inside the MCP. PduP/Q saturation could also be achieved with a very low cell membrane permeability to propionaldehyde, causing an accumulation of propionaldehyde in the cytosol, but at the cost of cytosolic aldehyde concentrations above the toxicity limit (Fig. S3). In addition, the membrane permeability to propionaldehyde required for this to occur (10^−7^ cm/s) is dramatically lower than a physiologically reasonable estimate (10^−3^ cm/s).

### MCPs function to enhance pathway flux

Decoupling PduP/Q saturation from cytosolic propionaldehyde concentration by encapsulation allows significantly greater carbon flux through the MCP metabolon than in the cases of enzyme scaffolding or no organization (Fig. 3C). Carbon flux per cell through PduP/Q in the MCP case (2.97*x*10^−13^ *μ*Mol/cell-s) is four orders of magnitude higher than in either the scaffold or no MCP cases (8.61*x*10^−17^ *μ*Mol/cell-s and 3.58*x*10^−17^ *μ*Mol/cell-s, respectively). Interestingly, this improvement is due solely to saturation of the PduP/Q enzymes; the flux through the PduCDE enzyme is similar with MCPs (6.49*x*10^−13^ *μ*Mol/cell-s) and without (7.41*x*10^−13^ *μ*Mol/cell-s). PduCDE production of aldehyde is sufficient in all four organizational cases, but without a substantial diffusion barrier in the form of the MCP membrane, the aldehyde leaks into the cytosol or the extracellular space before it can be utilized by the PduP/Q enzymes.

To quantitatively evaluate our model we estimate the growth rate resulting from the predicted flux through PduP/Q in the MCP case. Most parameter estimates were made from literature or a priori (Table 1); the nonspecific MCP membrane permeability *k*_*c*_ was set to the value that resulted in the greatest flux through the PduP/Q enzymes in our model (Fig. 2A). Our model predicts a flux of 2.97*x*10^−13^ *μ*mol/cell-s for a cell with MCPs, equivalent to 1.74*x*10^−5^ pg/cell-s, when the external 1,2-PD concentration is 55 mM. Approximately one-half of this flux can be used for cell growth, so assuming that a bacterial cell has a dry weight of approximately 0.3 pg [23], our model predicts a time of approximately 9 hours for a cell with MCPs to metabolize enough biomass through the Pdu MCP metabolon to accumulate the mass of one daughter cell (Fig. 2B). This value is in good agreement with experimentally measured doubling times for the growth of *Salmonella enterica* on 55 mM 1,2-PD of approximately 5-10 hours [21]. We believe the model is well suited to address batch-wise experimental results of this kind because a typical experiment measuring the growth of *Salmonella* on 1,2-PD takes place over tens of hours, while our model has characteristic timescales on the order of seconds, at maximum (Table 2). This allows us to treat a *Salmonella* growth experiment as being at a pseudo-steady state relative to our model, and accounts for the good congruence between our model predictions and the experimental measurements of steady-state *Salmonella* growth rates on 1,2-PD.

**Table 2.** Characteristic times for important processes in our model.

| Process | Representation | Characteristic time (s) |
| --- | --- | --- |
| Diffusion in the cell | $\tau_{cell}^{diff} = \frac{R_b^2}{D}$ | $2.5 \times 10^{-4}$ |
| Diffusion in the Pdu MCP | $\tau_{MCP}^{diff} = \frac{R_c^2}{D}$ | $1 \times 10^{-5}$ |
| Passive transport across the cell membrane | $\tau_{cell}^{trans} = \frac{R_c^2}{3k_m}$ | $4 \times 10^{-1}$ |
| Passive transport across the MCP membrane | $\tau_{MCP}^{trans} = \frac{R_c^2}{3k_c}$ | $3 \times 10^{-1}$ |
| PduCDE activity | $\tau_{CDE} = \frac{K_{MCDE}}{V_{CDE}}$ | $3 \times 10^{-3}$ |
| PduPQ activity | $\tau_{PQ} = \frac{K_{MPQ}}{2V_{PQ}}$ | $3 \times 10^{-1}$ |

Another putative MCP function is the prevention of aldehyde loss into the growth medium. We quantify this phenomenon in our model as the net flux of propionaldehyde across the cell membrane into the extracellular space (so-called “propionaldehyde leakage”). This leakage is lower with MCPs than without (3.52*x*10^−13^ *μ*Mol/cell-s as compared to 7.32*x*10^−13^ *μ*Mol/cell-s), but is significant relative to the flux through PduP/Q in either case when the external 1,2-PD concentration is 55 mM (Fig. 3D). For the case with MCPs, the flux through PduP/Q is approximately the same as the leakage flux, while leakage is over twice the PduP/Q flux in the no-MCP case. The flux through PduP/Q, the leakage flux, and the concentrations of propionaldehyde in the cytosol and the MCP, are plotted for a range of external 1,2-PD concentrations in Figure 4. Figure 4A shows the absolute metabolite concentrations in the MCP and cytosol in cells with and without MCPs and Figure 4B shows the absolute aldehyde leakage and PduP/Q flux for these two cases.

**Figure 4.**
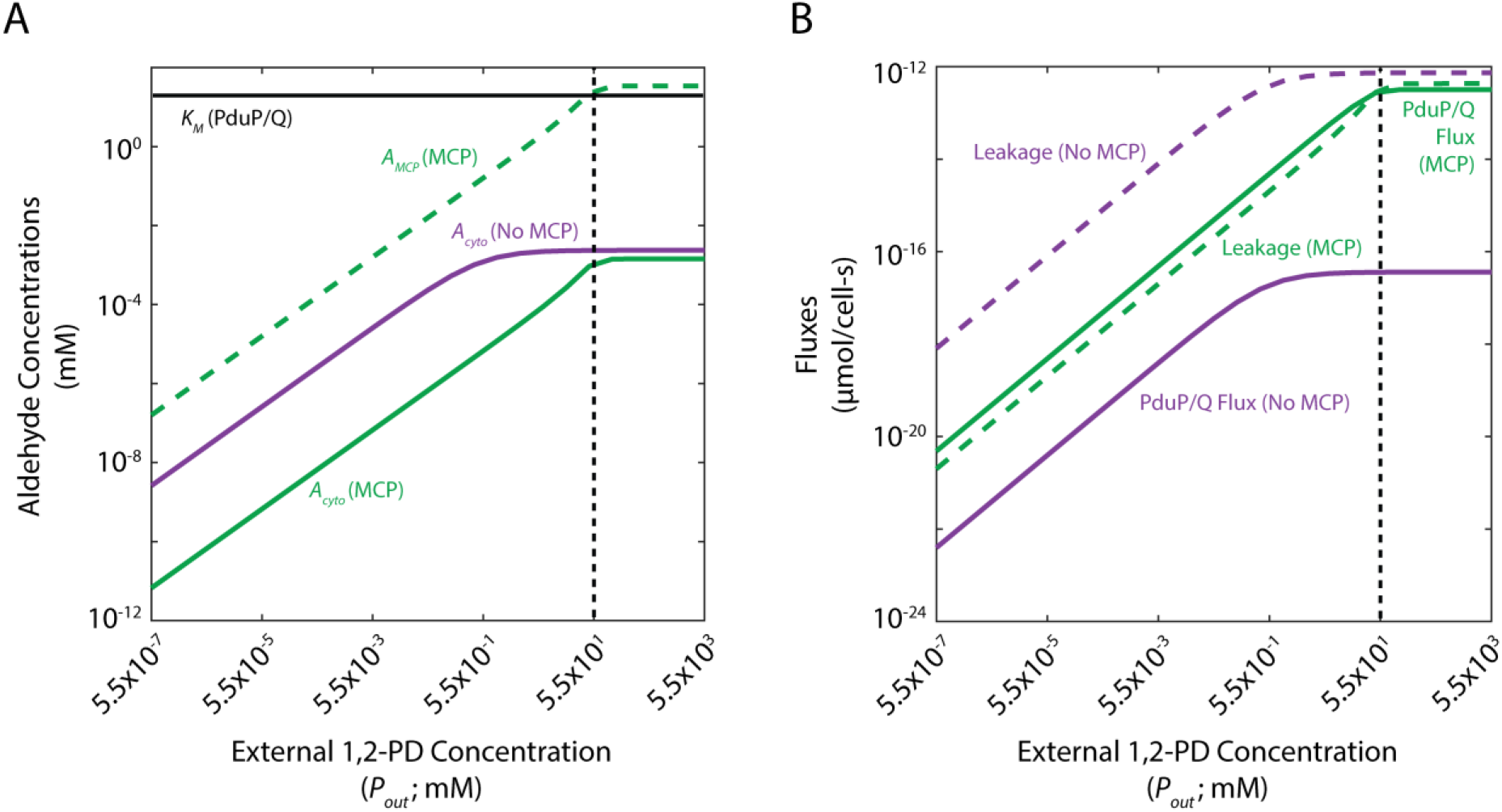
(A) Cytosolic propionaldehyde concentrations (*A_cyto_*) with and without MCPs and MCP propionaldehyde concentrations (*A_MCP_*) with MCPs and (B) steady-state fluxes through PduP/Q and propionaldehyde leakage across the cell membrane with and without MCPs. The baseline external 1,2-PD concentration is shown with a black dashed line; the *K_M_* of PduP/Q is shown in (A) with a black solid line.

At high external 1,2-PD concentrations, the cytosolic propionaldehyde concentration is comparable with and without MCPs (Fig. 4A). However, the kinetically relevant propionaldehyde concentration in the vicinity of the PduP/Q enzymes is much higher in the MCP when MCPs than in the cytosol without MCPs. The flux through the PduP/Q enzymes is therefore much higher and the MCP functions primarily for flux enhancement. On the other hand, at low external 1,2-PD concentrations, the cytosolic propionaldehyde concentration is lower with MCPs, resulting in reduced aldehyde leakage into the extracellular space. Conversely, the flux through the PduP/Q enzymes is more similar with and without MCPs in the case of low external 1,2-PD (and very low in either case), since the kinetically relevant propionaldehyde concentrations in the vicinity of the PduP/Q enzymes are more similar than at higher external 1,2-PD concentrations (Fig. 4B). In the case of low external 1,2-PD concentration, therefore, the relative difference in propionaldehyde leakage into the extracellular space is large, and the MCP functions primarily for aldehyde leakage reduction. It should also be noted that the flux through PduP/Q remains higher at all external 1,2-PD concentrations when the enzymes are localized to the MCP, even though the relative flux changes by an order of magnitude. There exists a transition from primarily flux enhancement to primarily aldehyde loss prevention as the external 1,2-PD concentration decreases. At high external 1,2-PD concentrations, cells with and without MCPs lose similar fluxes of propionaldehyde to the extracellular space, but cells with MCPs experience much greater flux through PduP/Q; this is due to saturation of the PduP/Q enzymes by the high propionaldehyde concentration inside MCPs. At low external 1,2-PD concentrations, on the other hand, cells with MCPs are more parsimonious with respect to propionaldehyde, but gain a smaller benefit in flux through PduP/Q since the PduP/Q enzymes are not saturated, even with MCPs (Fig. S4).

### Passive mechanisms are sufficient to support enzyme saturation at high external 1,2-PD concentrations, but active transport is necessary at low external 1,2-PD concentrations

We next determined the range of external 1,2-PD concentrations that saturate the PduP/Q enzymes in the MCP system. We explored this question using phase space representations of the saturation of the PduCDE and PduP/Q enzymes. In each phase space plot, two model parameters are varied and the effect on enzyme saturation is shown. We illustrate regions of parameter space in which neither enzyme is saturated (grey), only PduCDE is saturated (orange), or both enzymes are saturated (blue). Saturation of PduP/Q without saturation of PduCDE was not observed. Also shown in each phase space are isolines illustrating the parameter values for which the cytosolic concentration of propionaldehyde is 10 nM (0.001% of the toxicity limit) and 1 uM (0.1% of the toxicity limit), as well as dotted lines indicating the baseline parameter estimates used in the model. Recall that the toxicity limit for intracellular propionaldehyde is approximately 8 mM. Phase space representations of this kind are useful because they allow examination of the behavior of the system over a very wide range of parameter space, encompassing the entire range of physically reasonable values for each parameter in question.

In Figure 5A, for instance, the saturation of PduCDE and PduP/Q is examined as a function of the value of the nonspecific MCP membrane permeability *k*_*c*_ and the external 1,2-PD concentration *P*_*out*_. The blue region indicating saturation of both PduCDE and PduP/Q occurs only at high *P*_*out*_ values comparable to or higher than the baseline concentration. PduCDE alone can be saturated for a wider range of *P*_*out*_, as indicated by the extent of the orange region. Interestingly, for a broad range of *P*_*out*_ concentrations, neither enzyme can be saturated no matter the value of the nonspecific MCP permeability *k*_*c*_. We therefore conclude that PduCDE and PduP/Q can be saturated by adjusting *k*_*c*_ for a *P*_*out*_ greater than 50 mM, but for lower *P*_*out*_ concentrations no value of *k*_*c*_ achieves enzyme saturation. We expect that these lower concentrations are relevant for MCP-mediated metabolism *in vivo* because we observe that a *PPdu-gfp* reporter is activated for 1,2-PD concentrations as low as 55 *μ*M [24].

**Figure 5.**
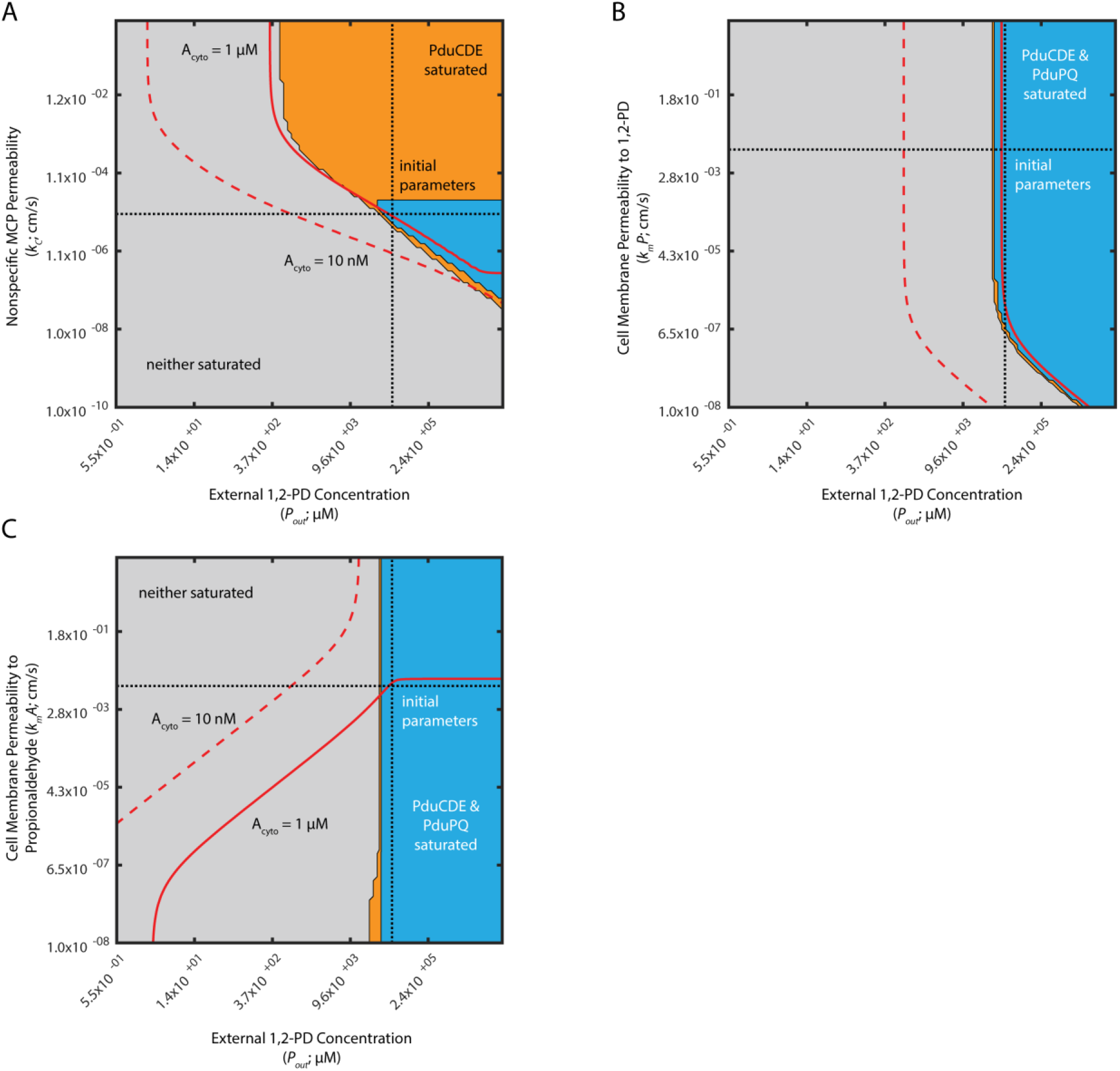
Saturation phase spaces of PduCDE and PduP/Q (A) with respect to *P_out_* and *k_c_*, (B) with respect to *P_out_* and 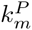, and (C) with respect to *P*_*out*_ and 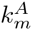. Regions of saturation (concentration of substrate > *K_M_* of the appropriate enzyme) are plotted in blue when PduCDE and PduP/Q are saturated, orange when only PduCDE is saturated, and in grey when neither enzyme is saturated. Red solid lines indicate when *A_cyto_* is 1 μM; red dashed lines indicate when *A_cyto_* is 10 nM. Black dashed lines indicate the baseline parameter values used in the model of the Pdu MCP.

Extending this analysis to the other passive diffusion mechanisms considered in the model, we find that PduP/Q can also be saturated for a range of external 1,2-PD concentrations by modulating the cell membrane permeability to 1,2-PD, 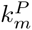, and the cell membrane permeability to propionaldehyde 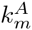 but that for each parameter there exists a lower limit of external 1,2-PD concentration (30 mM and 20 mM, respectively) below which PduP/Q cannot be saturated by passive mechanisms (Fig. 5B, C).

The *pduF* ORF of the *S. enterica* Pdu operon is a putative membrane protein, and is speculated to encode a 1,2-PD transporter [25]. We therefore explored the possible role of active 1,2-PD transport across the cell membrane in the saturation of the PduP/Q enzymes. Figure 6 shows phase space representations of PduCDE and PduP/Q saturation with respect to active 1,2-PD transport and two passive transport parameters when the external 1,2-PD concentration is 55 mM; Figure S5 shows the same analysis when the external 1,2-PD concentration is 0.5 mM. We find that active transport of 1,2-PD is dispensable at high external 1,2-PD concentrations (*i.e.* 55 mM), but not at lower external 1,2-PD concentrations (*i.e.* 0.5 mM). When the external 1,2-PD concentration is 55 mM, the nonspecific MCP membrane permeability, *k*_*c*_, can adopt a value such that both PduCDE and PduP/Q are saturated for any value of the velocity of active 1,2-PD transport across the cell membrane, *j_c_* (Fig. 6A). The same is true of the cell membrane permeability to 1,2-PD, 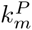, when the external 1,2-PD concentration is 55 mM (Fig. 6B). In contrast, when the external 1,2-PD concentration is 0.5 mM, there exists a minimum values (1 cm/s) of the velocity of active 1,2-PD transport across the cell membrane, *j_c_*, below which PduP/Q cannot be saturated solely by modulating the MCP shell passive diffusion parameter *k*_*c*_ (Fig. S5). Similar minima exist at active transport velocities of 2*x*10^−5^ cm/s and 0.3 cm/s for the cell membrane passive diffusion parameters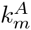 and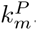. At this lower external concentration of 1,2-PD, therefore, active transport of 1,2-PD across the cell membrane may play an important role. Indeed, with high rates of active transport of 6*x*10^3^ cm/s, PduP/Q can be saturated for an extremely wide range of external 1,2-PD concentrations (Fig. S6).

**Figure 6.**
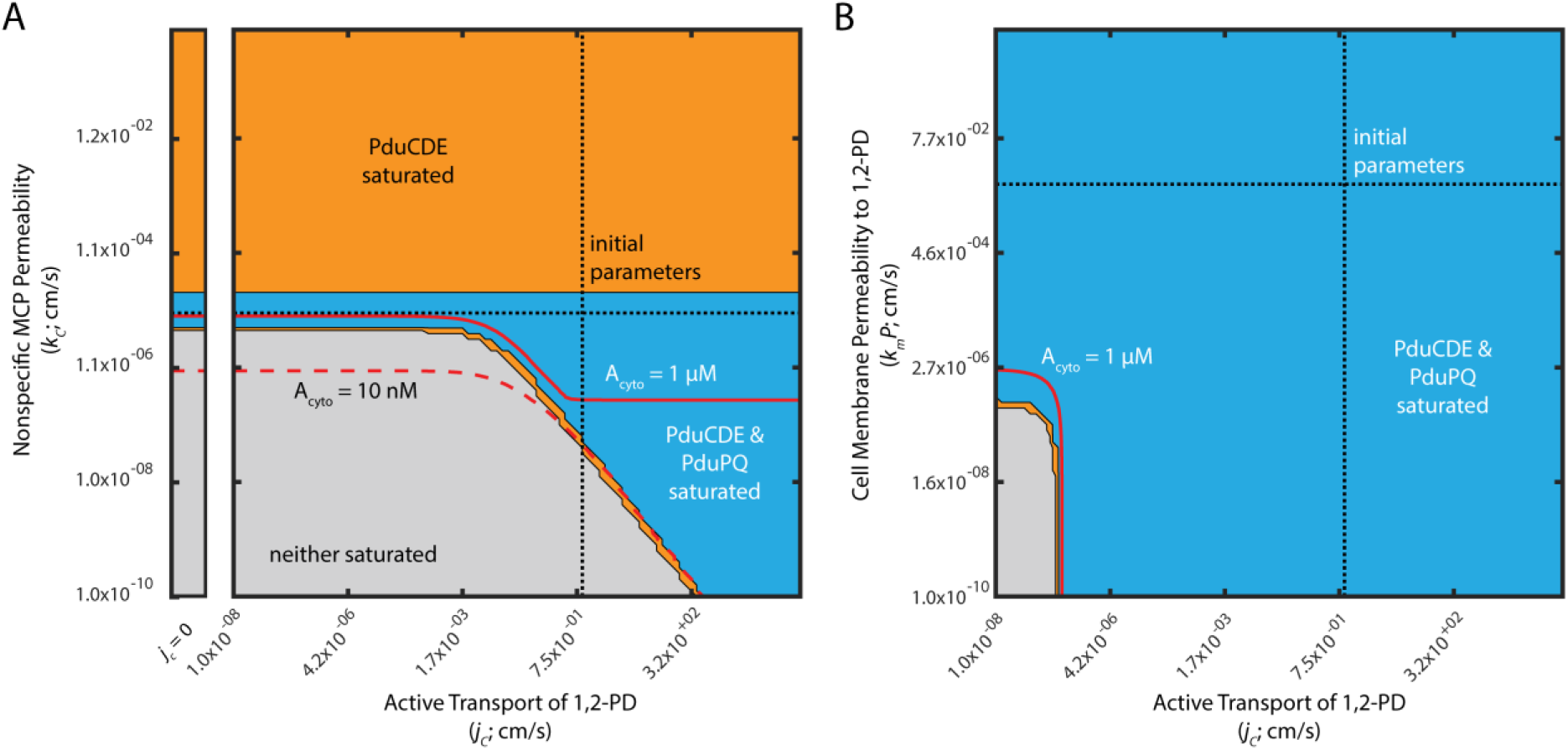
Saturation phase spaces of PduCDE and PduP/Q with respect to (A) *j_c_* and k_c_ and (B) with respect to *j_c_* and 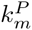 when Pout is 55 mM. Regions of saturation (concentration of substrate > K_M_ of the appropriate enzyme) are plotted in blue when PduCDE and PduP/Q are saturated, orange when only PduCDE is saturated, and in grey when neither enzyme is saturated. Red solid lines indicate when *A_cyto_* is 1 μM; red dashed lines indicate when *A*_*cyto*_ is 10 nM. Black dashed lines indicate the baseline parameter values used in the model of the Pdu MCP. Inset to the left in (A) reflects the behavior when j_c_ = 0.

We can further understand these trends by examining the analytical solution to the model. The relative contribution of active transport to 1,2-PD transport across the cell membrane (as compared to passive diffusion) is expressed by the quantity 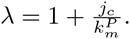 This suggests that active transport can only be significant if 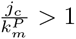. Furthermore, transport of 1,2-PD across the cell membrane only impacts the steady-state 1,2-PD concentration in the MCP when λp∗ ≈ Г_C DE_ E, where *λp* ^*∗*^ represents active and passive transport across the cell membrane and ГC DE represents the balance between transport and reactions processes. Therefore, active transport only bears on the solution when 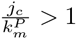 and λp∗ ≈ Г_C DE_ E. In Figure 6B, when the external 1,2-PD concentration is high, λp∗ is large relative to Г_C DE_ and changes in λ are inconsequential except at large λ values (when 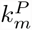 is small). When the external 1,2-PD concentration is low, however, as in Figure S4, λp∗ ≈ Г_C DE_ E and active transport is therefore consequential for a large range of 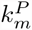. We also observe that active transport only impacts saturation when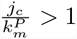, as expected from the analytical solution.

### Selective MCP permeability is not absolutely required to saturate PduP/Q

Experimental results suggest that the protein membrane surrounding the Pdu MCP might exhibit selective permeability [26]. We therefore explored under what conditions such selective permeability was advantageous for MCP function. We first consider the simple case of nonspecific permeability.

We find that an optimal non-selective MCP shell permeability exists with respect to the kinetically relevan
t propionaldehyde concentration in the vicinity of the PduP/Q enzymes (Fig. 7A). This optimum value (10^−5^ cm/s) of a single nonspecific permeability *k*_*c*_ reflects a tradeoff between 1,2-PD entry to the MCP and trapping of propionaldehyde within the MCP. The non-selective permeability must be high enough for adequate entry of the PduCDE substrate 1,2-PD, but low enough to contribute to the accumulation of the PduP/Q substrate propionaldehyde within the organelle. This can be seen clearly when the permeabilities of the MCP membrane to 1,2-PD and propionaldehyde are varied separately (Fig. 7B,C). Lower permeability to propionaldehyde is unambiguously advantageous for trapping propionaldehyde in the MCP. Higher permeability to 1,2-PD, 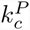, also increases propionaldehyde concentration inside the MCP, until the permeability is sufficient to equalize the cytosolic and MCP 1,2-PD concentrations, at which point there is no further improvement.

**Figure 7.**
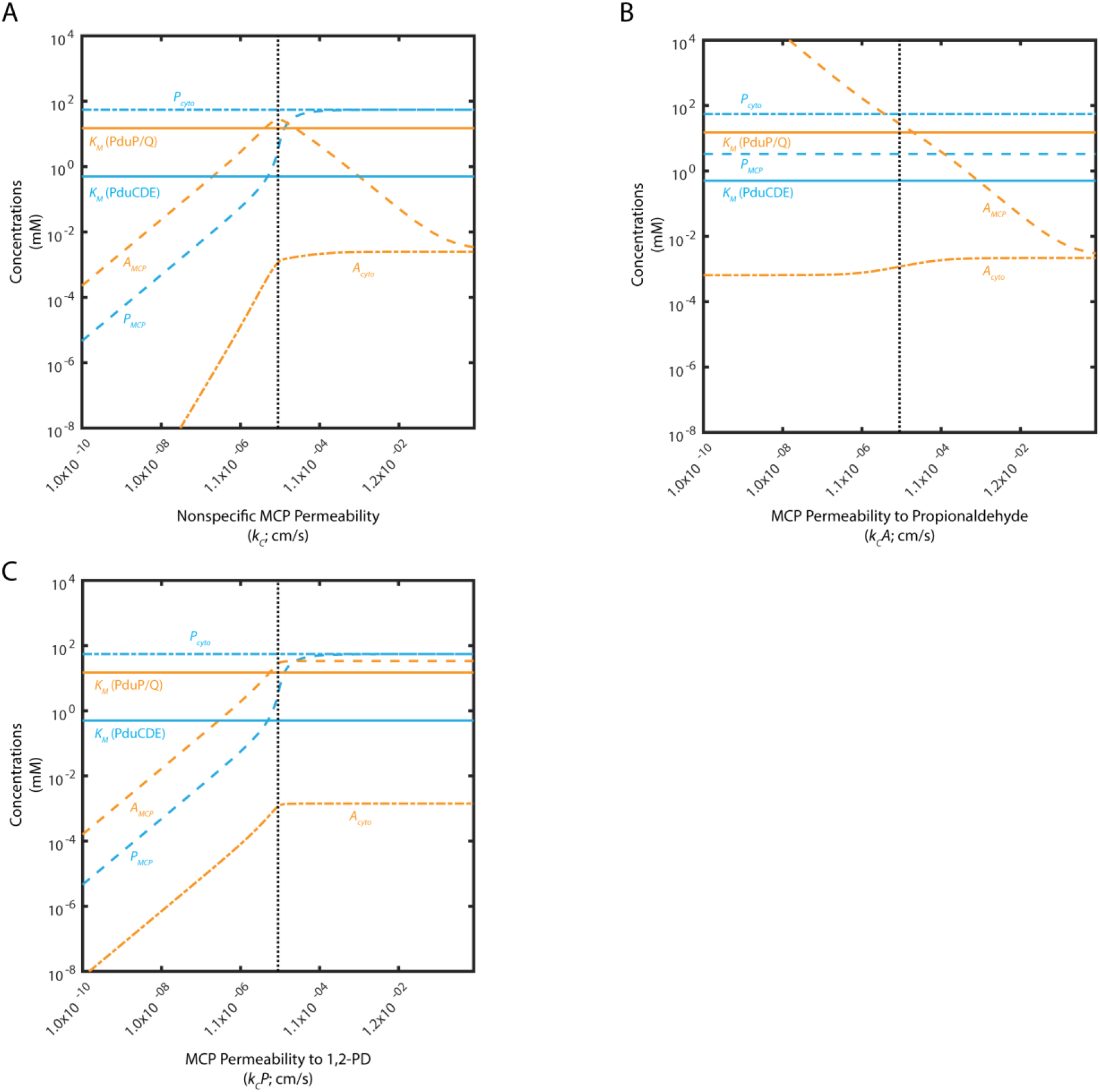
Mean concentrations of 1,2-PD and propionaldehyde in the MCP (P*_MCP_*; A*_MCP_*) and cytosol (*P*_*cyto*_; A_cyto_) as a function of (A) *k_c_* when 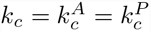; (B) 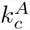; and (C) 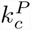. *K*_*M*_ of PduCDE and PduP/Q are shown as solid lines. The baseline permeabilities are shown with a black dashed line.

The presence of an optimal permeability persists when the ratio of 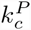 to 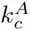 is fixed at 0.1 or 10 and the values are varied together, maintaining this ratio (Fig. S7). Moreover, decreasing *k*_*c*_ entails a tradeoff between leakage prevention and flux enhancement, the two aspects of Pdu MCP function (Fig. S8). At low *k*_*c*_ (less than 10^−7^cm/s), aldehyde leakage is prevented, but flux is low; near the optimal *kc* for flux enhancement (10^−5^cm/s), leakage is comparable with and without MCPs. At high *kc* (greater than 1), the system approaches the case of scaffolding, with only a small flux enhancement and no leakage prevention.

We determine the potential benefit of selective permeability by examining the enzyme saturation phase space with respect to the specific permeabilities 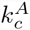 and 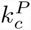 (Fig. 8). The 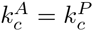 line, along the diagonal, in this subspace indicates the c c performance of the system when the MCP permeability is non-selective. This line passes through the blue region in which PduP/Q is saturated, indicating that selective permeability is not absolutely required for efficient performance. However, selective permeability permits a broader range of 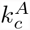 and 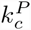 values to saturate PduP (Fig. 8). It is interesting to note that MCP permeability to propionaldehyde must be lower than or equal to MCP permeability to 1,2-PD. As observed above, decreasing MCP permeability to propionaldehyde is unambiguously beneficial for flux, while decreasing the MCP permeability to 1,2-PD below the MCP permeability to propionaldehyde is detrimental to relative flux (Fig. S8). It is also important to note, however, that increasing the concentration of propionaldehyde inside the MCP beyond the concentration required to saturate PduP/Q is not beneficial, since there is no increase in flux, but propionaldehyde leakage increases.

**Figure 8.**
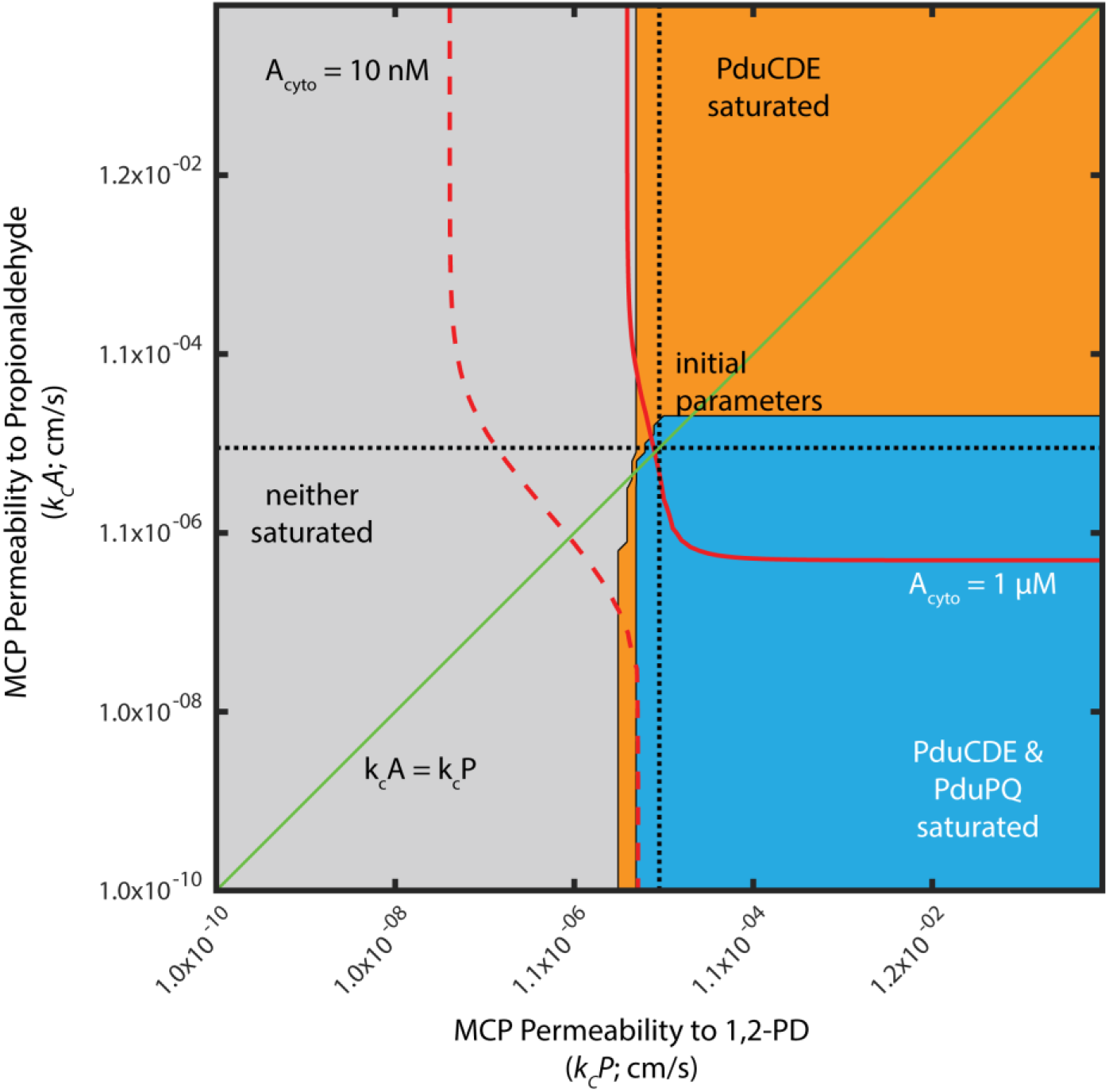
Saturation phase space of PduCDE and PduP/Q with respect to 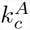 and 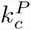. Regions of saturation (concentration of substrate > K_M_ of the appropriate enzyme) are plotted in blue when PduCDE and PduP/Q are saturated, orange when only PduCDE is saturated, and in grey when neither enzyme is saturated. Red solid lines indicate when *A*_*cyto*_ is 1μM; red dashed lines indicate when A_cyto_ is 10 nM. Black dashed lines indicate the baseline parameter values used in the model of the Pdu MCP. Green line indicates 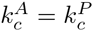.

## Discussion

By analyzing the closed-form analytical solution to our mechanistic model of Pdu MCP function, we first find that flux enhancement may play a more significant role in MCP function than previously thought. Secondly, our results suggest that active transport of 1,2-PD across the cell membrane may in many cases contribute significantly to MCP function. Lastly, we find that, while not always required for MCP function, selective MCP membrane permeability can enhance MCP function.

### Flux enhancement is a key facet of MCP function

Experimental investigations of Pdu MCP function have consistently demonstrated two key phenotypes for strains that express the Pdu enzymes but fail to form MCPs: a slower growth rate and increased propionaldehyde concentration in the media [21]. Slower growth could be attributable to two phenomena: “passive” growth retardation due to lower carbon flux through the Pdu metabolon, and “active” growth retardation due to the toxic effects of propionaldehyde in the cytosol. This second form of growth defect is linked to the accompanying observation that strains lacking MCPs have an increased rate of propionaldehyde leakage into the extracellular space. These two forms of growth retardation cannot be distinguished by an *in vivo* experiment measuring the growth of cells and the bulk concentrations of the various metabolites. Our simple model supports the idea that both of these phenotypes contribute to changes in growth rate: we observe that cells with MCPs have both higher flux through PduP/Q and lower cytosolic concentrations of propionaldehyde than cells without MCPs, and that cells with MCPs exhibit lower aldehyde flux into the growth medium than cells without MCPs. Additionally, experiments indicate similar 1,2-PD depletion from the growth medium with and without MCPs, a phenotype that is evident in the comparable PduCDE fluxes predicted by our model with and without MCPs [21]. Together these observations indicate that our model captures the important principles of Pdu MCP function. Indeed, we propose that while toxicity reduction likely plays a role in MCP function, flux enhancement is a crucial, and underappreciated, consequence of encapsulation. This in turn suggests that experiments should explore the relative contributions of flux enhancement and toxicity mitigation in more detail. For instance, the effects of propionaldehyde toxicity could be characterized by determining the degree to which aldehyde leakage from MCP-defective cells leads to increased formation of covalent adducts to the cellular genome and proteome.

### MCP function depends on extrinsic factors

Many behaviors in our model depend strongly on external 1,2-PD concentration. Therefore, care must be taken in applying results from experiments conducted at high 1,2-PD concentrations (55 mM) in the laboratory to Pdu MCP function inside the host, and in interpreting the results of batch-wise culture experiments in which the external 1,2-PD concentration changes during the experiment, decreasing from 55 mM to 1 mM [21]. We find that MCP function, as quantified by the relative PduP/Q flux and the relative propionaldehyde flux into the extracellular space, changes in response to factors extrinsic to the cell, such as external 1,2-PD concentration. Indeed, these may change during a single experiment: for example, cells without MCPs may be observed to leak slightly more propionaldehyde into the growth medium than cells with MCPs when grown at high external 1,2-PD concentrations, but this leakage discrepancy may increase over the course of a batch experiment during which 1,2-PD is depleted from the growth medium. Whether or not PduP/Q can be saturated without active transport also depends on external 1,2-PD concentration; our model suggests that active transport of 1,2-PD across the cell membrane is dispensable at high external 1,2-PD concentrations, but not as the external 1,2-PD concentration decreases. These mechanistic observations suggest that experiments to determine the function of the *pduF* gene product, for instance, should be undertaken in low 1,2-PD concentration so that active transport is made relevant to growth.

It is unknown what external concentration of propionaldehyde is encountered in the host gut by invading *Salmonella*, but the concentration is likely to be low due to the toxicity of aldehyde species to eukarya. With respect to laboratory growth experiments, the propionaldehyde concentration in the growth medium is observed to increase in batch-wise growth experiments, up to concentrations around 10 mM [21]. The calculations above were performed assuming no external propionaldehyde, but we tested our model at a range of external propionaldehyde concentrations (1 *μ*M, 1 mM, and 10 mM; Fig. S9) and found that the qualitative behavior of the system did not change with increasing external propionaldehyde concentration. The regions of parameter space in which PduP/Q could be saturated were broader at higher propionaldehyde concentrations, but at the cost of high cytosolic aldehyde concentrations equilibrated with the external concentration.

### Selective MCP membrane permeability is not required, but is often advantageous

Recent evidence suggests that the Pdu MCP shell may be selectively permeable to propionaldehyde as compared to other metabolites [26]. Our model suggests that, while the saturation of PduP/Q enzymes can be achieved without selective permeability in particular cases, it is true in general that selective permeability can enhance Pdu MCP function. In contrast, selective permeability of the MCP shell does not improve the function of a related system, the carbon concentrating mechanism [22]. The monotonic benefit of selective permeability in the Pdu system is due to PduCDE catalyzing an essentially irreversible reaction, whereas the enzyme playing the equivalent role in the carbon concentrating mechanism is reversible. Trapping of propionaldehyde in the MCP by decreasing the relative permeability of the MCP shell to propionaldehyde can therefore saturate the PduP/Q enzymes if the permeability of the MCP to 1,2-PD is sufficiently high. Similarly, if the permeability of the MCP to propionaldehyde is sufficiently low, the permeability to 1,2-PD can be increased to saturate PduP/Q. This putative trapping mechanism is in congruence with the *in vitro* observation that small molecule efflux from a protein nanoreactor can be affected by the chemical character of the reactor pores [27]. Studies of substrate channeling in enzyme scaffolds also emphasize the importance of creating a high substrate concentration in the vicinity of downstream enzymes [28,29]. In the case of a microcompartment, the shell diffusion barrier creates a concentration differential across the MCP shell, rather than a very high local concentration of enzymes enforcing a local diffusion gradient. Interestingly, and contrary to previous studies, our results suggest that selective permeability, while often advantageous, is by no means required for significant flux enhancement (Fig. 3). Further experimental determination of the MCP membrane permeability is required to determine in what parameter regime the Pdu MCP system operates, and whether selectivity is required in vivo. Moreover, since our model predicts an optimal nonspecific Pdu MCP permeability of 10^−5^ cm/s, we can compare this value to future experimental data for the various relevant metabolites.

### Outstanding questions for modeling and experiment

Our results have important implications for microbiological studies of MCP function and for the application of encapsulation to synthetic biology. Interestingly, our model indicates that the encapsulation of enzymes in a subcellular compartment can dramatically improve flux through the encapsulated pathway, in addition to reducing the concentrations of intermediates in the cytosol, simply by imposing a non-specific diffusion barrier. This reinforces the notion that encapsulation is a promising strategy to improve the yield and titer of heterologous enzymatic pathways that fail to function in a cytosolic context. However, further investigation is required to inform the selection of appropriate pathways for encapsulation. We also find that, for the Pdu system, encapsulation is superior to scaffolding in enhancing pathway flux, and future efforts will explore what characteristics render certain pathways amenable to encapsulation or scaffolding.

Many questions also remain to be addressed both computationally and experimentally with respect to native MCP function. First, in our model we neglect NAD+ /NADH cofactor recycling at steady state by setting the rate of PduP catalysis equal to that of PduQ. Future models could incorporate cofactor-dependent kinetics of PduP/Q, in addition to including the effect of *B*_12_ recycling within the MCP shell on the kinetics of PduCDE. Also lacking at this time are detailed simulations of the structure and dynamics of the Pdu membrane pores, particularly with respect to the diffusion of species of varying size and charge. High resolution modeling, such as molecular dynamics simulations of the pore, could aid in determining how low permeabilities of 10^−5^ cm/s could be achieved. Studies of this kind would be complemented by direct experimental measurements of the permeability of the MCP membrane to various species. Our model makes quantitative predictions for the permeabilities of small molecule metabolites such as 1,2-PD and propionaldehyde, and also for the permeability of the MCP shell to cofactors like NAD^+^ /NADH (insomuch as we assume that the MCP shell is impermeable to these species).

Also of interest are experimental investigations as to the possibility of active 1,2-PD transport across the cell membrane. Our results suggest that such active transport would be advantageous at low external 1,2-PD concentrations but dispensable at high external 1,2-PD concentrations, complicating experiments. The putative membrane-bound Pdu gene product *pduF* is of primary interest in this regard. Lastly, experimental observations of the absolute 1,2-PD and propionaldehyde concentrations encountered by invading *Salmonella* in the host gut and in the environment at large would be valuable in constraining future modeling efforts to pathogenically relevant metabolite concentrations, and in comparing the conditions faced by free-living and host-associated pathogens.

## Supporting Information

### S1 Fig

**Comparison of analytical solution assuming constant concentrations in the MCP (solid lines) and numerical solutions from the edge (circles) and center (triangles) of the MCP for 1,2-PD (blue) and propionaldehyde (orange).** The baseline parameter values are shown with a black dashed line. The K_M_ of the PduCDE and PduP/Q enzymes are plotted in blue and orange lines, respectively.

### S2 Fig

**Concentration profiles as a function of r for a cell with (A) no MCPs; (B) a scaffold with no diffusion limitation** (*k*_*c*_ = 10^3^); **(C) MCPs** (*k*_*c*_ = 10^−5^); and **(D) sparingly permeable MCPs** (*k*_*c*_ = 10^−7^). 1,2-PD in the MCP (*P*_*MCP*_) and in the cytosol (*P*_*cyto*_) are plotted in blue and propionaldehyde in the MCP (*A*_*MCP*_) and in the cytosol (*A*_*cyto*_) in orange. The *K*_*M*_ of the PduCDE and PduP/Q enzymes are plotted in blue and orange dashed lines, respectively.

### S3 Fig

**(A) Cytosolic aldehyde concentration** (*A*_*cyto*_ **) with and without MCPs and MCP aldehyde concentration (***A_MCP_*) **with MCPs; (B) relative carbon flux through PduP/Q 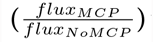 and relative aldehyde leakage rate 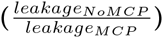; (C) relative flux through the PduP/Q enzymes (with MCPs/without MCPs) and relative propionaldehyde leakage across the cell membrane (without MCPs/with MCPs) as a function of 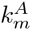.** The baseline 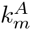 value is shown with a black dashed line.

### S4 Fig

**Relative flux through the PduP/Q enzymes (with MCPs/without MCPs) and relative propionaldehyde leakage across the cell membrane (without MCPs/with MCPs) as a function of external 1,2-PD concentration.** The baseline external 1,2-PD concentration is shown with a black dashed line.

### S5 Fig

**Saturation phase spaces of PduCDE and PduP/Q with respect to (A)***j*_*c*_ and *k*_*c*_, **(B) with respect to***j*_*c*_ and 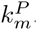 **, and (C) with respect to** *j*_*c*_ and 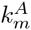 **when *P*_*out*_ is 0.5 mM**. Regions of saturation (concentration of substrate > K_M_ of the appropriate enzyme) are plotted in blue when both enzymes are saturate, orange when only PduCDE is saturated, and in grey when neither enzyme is saturated. Red solid lines indicate when *A*_*cyto*_ is 1 *μ*M; red dashed lines indicate when *A*_*cyto*_ is 10 nM. Black dashed lines indicate the baseline parameter values used in the model of the Pdu MCP.

### S6 Fig

**Saturation phase spaces of PduCDE and PduP/Q with respect to (A)** *P*_*out*_ **and** *j*_*c*_ **, and (B) with respect to** *P*_*out*_ **and** 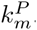. Regions of saturation (concentration of substrate > *K*_*M*_ of the appropriate enzyme) are plotted in blue when both enzymes are saturate, orange when only PduCDE is saturated, and in grey when neither enzyme is saturated. Red solid lines indicate when *A*_*cyto*_ is 1 *μ*M; red dashed lines indicate *A*_*cyto*_ is 10 nM. Black dashed lines indicate the baseline parameter values used in the model of the Pdu MCP.

### S7 Fig

**Mean concentrations of 1,2-PD and propionaldehyde in the MCP (*P*_*MCP*_; *A*_*MCP*_) and cytosol (*P*_*cyto*_; *A*_*cyto*_) as a function of (A) 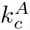 when 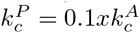 and (B) 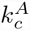 when** 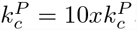. *K*_*M*_ of PduCDE and PduP/Q are shown as solid lines. The baseline permeabilities are shown with a black dashed line.

### S8 Fig

**(A,C,E) Cytosolic aldehyde concentration** (*A*_*cyto*_) **with and without MCPs and MCP aldehyde concentration** (*A*_*cyto*_) **with MCPs; (B,D,F) relative carbon flux through PduP/Q (fluxMCP/fluxNoMCP) and relative aldehyde leakage rate (leakageNoMCP/leakageMCP) as a function of (A,B)** 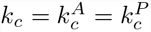; (**C,D**) 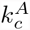; and (**E,F**) 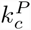. The baseline permeabilities are shown with a black dashed line.

### S9 Fig

**(A, C, E) Saturation phase space of PduCDE and PduP/Q with respect to** 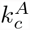 **and 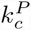 and (B, D, F) saturation phase space of PduCDE and PduP/Q with respect to** 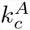 **and** *j*_*c*_ **for external propionaldehyde concentrations of (A, B) 1** *μ* **M, (C, D) 1 mM, and (E, F) 10 mM.** Regions of saturation (concentration of substrate > *K*_*M*_ of the appropriate enzyme) are plotted in blue when both enzymes are saturated, orange when only PduCDE is saturated, and in grey when neither enzyme is saturated. Red solid lines indicate when *A*_*cyto*_ is 1 *μ*M; red dashed lines indicate when *A*_*cyto*_ is 10 nM. Black dashed lines indicate the baseline parameter values used in the model of the Pdu MCP. Green line in (A, C, E) indicates when 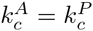.

## Acknowledgments

The authors wish to thank the Tullman-Ercek and Savage Labs for stimulating discussions, and Avi Flamholz for his efforts in building the CCM model on which this work is based.

## Appendix A

We can find the following complete analytical solutions in the cytosol as a function of the concentrations in the MCP

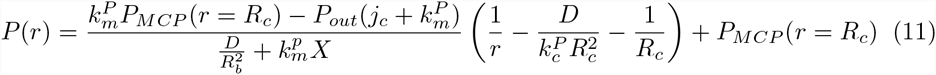

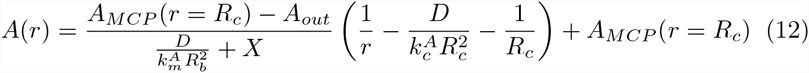

Where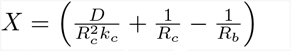

We can then use the solution in the cytosol to generate the following boundary condition at the MCP membrane:

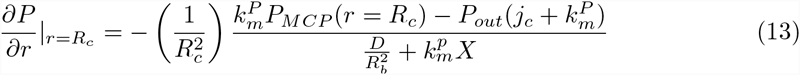

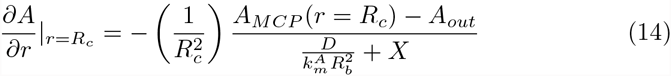

First, consider the mass balance on *A*_*M C P*_:

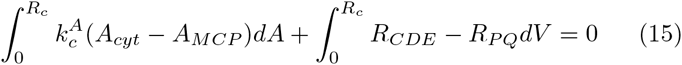

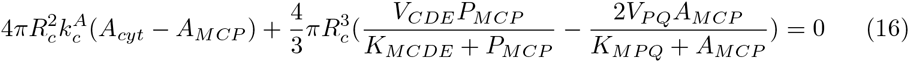

And similarly for *P*_*M C P*_:

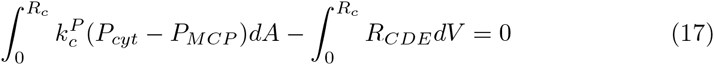

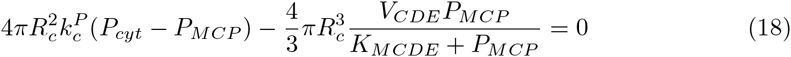

We now assume that the concentrations in the MCP are constant since *ξ >>* 1. First we solve for *P*_*MCP*_, as this does not depend on *A*_*MCP*_ due to the irreversibility of PduCDE. We simplify the solution by defining the following important timescales, assuming that 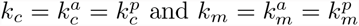(Table 2):

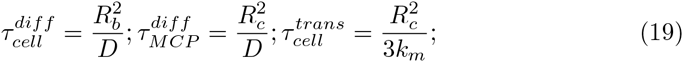

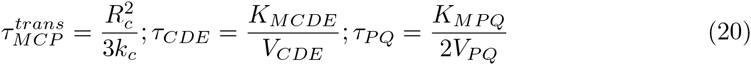

Letting 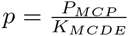, 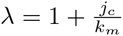, 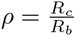 and 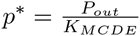 the solution for *p* is therefore as follows:

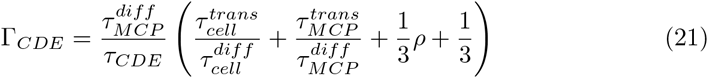

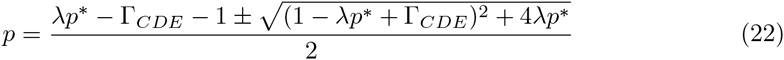

Furthermore, if PduCDE is saturated,

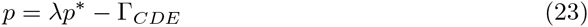

We can estimate the magnitudes of these various timescales based on the baseline model parameters (Table 2) and thence analyze the magnitude of the various terms in Г_*C DE*_.

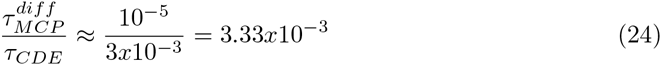

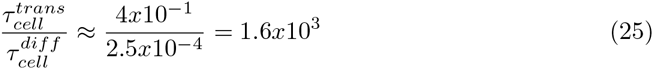

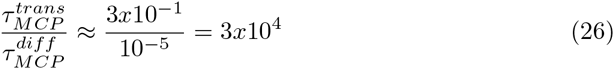

Therefore, for the baseline model parameter values in Table 1,

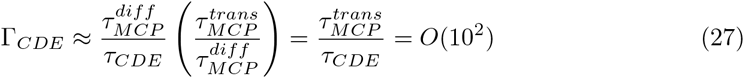

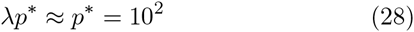

Suggesting that in the vicinity of the baseline parameter values, the solution for *P* in the MCP is governed by the relative timescales of the transport of 1,2-PD in and out of the MCP and the reaction of 1,2-PD to propionaldehyde by PduCDE, as well as by the external 1,2-PD concentration.

Now we can find *A_MCP_* similarly, given the solution for *P*_*M CP*_. Letting 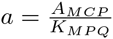, 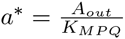 and 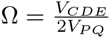

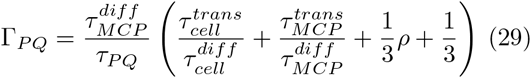

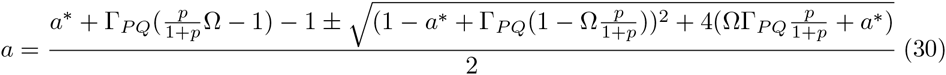

If PduCDE is saturated and *A*_*out*_ is negligible, then

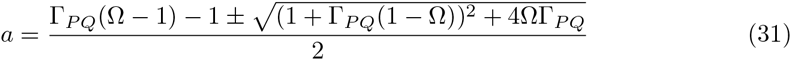

And if both PduCDE and PduPQ are saturated and *A*_*out*_ is negligible, then

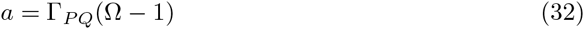

We can analyze the relative magnitudes of the timescales in Г_*PQ*_ as above, assuming the baseline parameter values in Table 1, and we find that

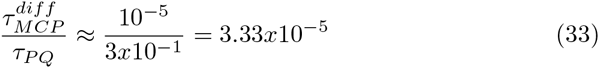

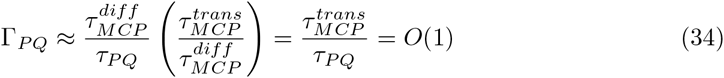

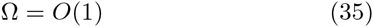

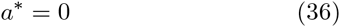

Suggesting that in the vicinity of the baseline parameter values, the solution for *A* in the MCP is governed by the relative timescales of the transport of propionaldehyde in and out of the MCP and the reaction of propionaldehyde by PduP/Q, as well as by the relative rates of PduCDE and PduP/Q.

Again, the solutions in the cytosol follow directly from these MCP solutions.

